# The gut microbiome and resistome of conventionally- vs. pasture-raised pigs

**DOI:** 10.1101/2023.03.02.530897

**Authors:** Devin B. Holman, Katherine E. Gzyl, Arun Kommadath

## Abstract

Conventional swine production typically houses pigs indoors and in large groups, whereas pasture-raised pigs are reared outdoors in lower stocking densities. Pigs in both production systems are usually fed a grain-based diet but pasture-raised pigs may also consume plants and soil. Antimicrobial use also differs with conventionally-raised pigs often being exposed to antimicrobials directly or indirectly to control and prevent infectious disease. However, antimicrobial use can be associated with the development and persistence of antimicrobial resistance. In this study, we used shotgun metagenomic sequencing to compare the gut microbiomes and resistomes of pigs raised indoors on a conventional farm with those raised outdoors on pasture. The microbial compositions as well as the resistomes of both groups of pigs were significantly different from each other. Bacterial species such as *Intestinibaculum porci, Pseudoscardovia radai*, and *Sharpea azabuensis* were relatively more abundant in the gut microbiomes of pasture-raised pigs and *Hallella faecis* and *Limosilactobacillus reuteri* in the conventionally-raised swine. The abundance of antimicrobial resistance genes (ARGs) was significantly higher in the conventionally-raised pigs for nearly all antimicrobial classes, including aminoglycosides, beta-lactams, macrolides-lincosamides-streptogramin B, and tetracyclines. Functionally, the gut microbiomes of the two group of pigs also differed significantly based on their CAZyme profiles, with certain CAZyme families associated with host mucin degradation enriched in the conventional pig microbiomes. We also recovered 1,043 dereplicated strain-level metagenome-assembled genomes (≥ 90% completeness and <5% contamination) to provide taxonomic context for specific ARGs and metabolic functions. Overall, the study provides insights into the differences between the gut microbiomes and resistomes of pigs raised under two very different production systems.

## INTRODUCTION

Pigs raised on conventional swine farms are typically housed indoors in large numbers under tightly controlled environmental conditions. Conversely, pasture-raised pigs are usually kept outdoors in smaller numbers where they have access to the soil and plants growing in the pasture. Antimicrobial use also often differs between these two production systems. Unlike conventionally-raised pigs, pasture-raised pigs are much more likely to be treated individually if required rather than as a group if an animal becomes ill. Antimicrobial use in food-producing animals such as swine has come under increased scrutiny due to its potential association with antimicrobial resistance [1]. However, even in the absence of antimicrobial exposure conventionally-raised or antimicrobial-free pigs raised indoors frequently still have a relatively diverse and rich resistome [2-4]. The resistome refers to antimicrobial resistance genes (ARGs) in all bacteria within a given environment and includes both acquired and intrinsic ARGs [5].

The gut microbiome plays an important role in the health and performance of swine. For example, the gut microbiome encodes many genes involved in the metabolism of nondigestible dietary carbohydrates into metabolites such as short-chain fatty acids (SCFAs) that can be used as an energy source by the host [6]. The swine gut microbiome is largely shaped by diet but is also influenced by age, antimicrobial use, disease state, environment, and genetics [4, 7, 8]. Cultivation of many microorganisms found in the mammalian gut remains challenging due to their often unknown growth requirements; however, techniques such as shotgun metagenomic sequencing can provide new insights into both the function and composition of microbiomes without the need for extensive culturing.

Pigs raised outdoors on pasture and without prior exposure to antimicrobials can provide important insights into how management practices can influence the swine gut microbiome and resistome. Therefore, our objective was to characterize the gut microbiomes and resistomes of commercial pigs that had been raised entirely outdoors without antimicrobial exposure and to compare them with conventionally-raised pigs that also did not receive antimicrobials.

## METHODS

### Animals and sampling

The pasture-raised pigs were crossbred Duroc, Tamworth, Berkshire, and Large Black breeds located on a commercial operation in Alberta where they were housed outdoors year-round and did not receive antimicrobials. Fecal samples were collected from pigs in various growth stages: sows (n =13), nursery (n =10), and growing/finishing (n = 6). Fecal samples were also taken from pigs at the swine unit of the Lacombe Research and Development Centre that were in similar growth stages and been weaned at 21 days of age. Animals were cared for in agreement with the Canadian Council for Animal Care (2009) guidelines. The Lacombe Research and Development Centre Animal Care Committee reviewed and approved all procedures and protocols involving animals.

### Metagenomic DNA extraction and sequencing

Metagenomic DNA was extracted from fecal samples using the QIAamp BiOstic bacteremia DNA kit (Qiagen, Mississauga, ON, Canada) as previously described [4]. DNA was quantified using the Quant-iT PicoGreen dsDNA Assay Kit (Thermo Fisher Scientific, Inc., Mississauga, Ontario, Canada) and metagenomic libraries were prepared using 50 ng of DNA and the NEBNext Ultra II DNA Library Prep Kit (New England BioLabs, Ipswich, MA, USA) as per the manufacturer’s recommendations. Size selection and purification of the metagenomic libraries was carried out using SparQ PureMag beads (Quantabio, Beverly, MA, USA) and an Illumina Genome Analyzer (Illumina Inc., San Diego, CA, USA) with a KABA SYBR Fast Universal qPCR Kit (Kapa Biosystems, Wilmington, MA, USA) was used for quantification. The metagenomic libraries were normalized and pooled and 225 pM was sequenced together with PhiX (1%) on a NovaSeq 6000 with a S4 flow cell (300 cycles) as per the manufacturer’s instructions. DNA from a mock community of 20 bacterial strains (MSA-2002; ATTC) was also extracted and sequenced as a positive control.

### Metagenomic sequencing analysis

Low quality reads and sequencing adapters were removed with fastp v.0.23.2 [9] using a 4-bp sliding window and a quality threshold of 15. Any reads with less than 100 bp were removed. Swine host and PhiX sequences were removed using Bowtie2 v.2.4.5 [10] to align the reads to three swine genome assemblies (Sscrofa11.1 [Duroc], Berkshire_v1 [Berkshire], USMARCv1.0 [Duroc x Landrace x Yorkshire]) and the *Escherichia* phage phiX174 genome (NC_001422) for removal. SAMtools v.1.14 [11] and BEDtools v.2.30.0 [12] were used to extract the reads that did not align to the swine and PhiX genomes and these paired fastq reads were used for all subsequent steps. Taxonomic classification of the metagenomic samples was carried out using Kaiju v. 1.9.0 (Menzel, Ng et al. 2016) and the NCBI non-redundant protein database (March 19, 2022). The unassembled reads were also screened for antimicrobial resistance genes (ARGs) using the resistance gene identifier (RGI) v. 6.0.1 and the Comprehensive Antibiotic Resistance Database (CARD) v.3.2.5 [13] with KMA [14] used for alignment. Carbohydrate-active enzyme (CAZyme) profiles were characterized by mapping the reads to the dbCAN2 (Zhang, Yohe et al. 2018) v. 09242021 using DIAMOND v. 2.0.14 (Buchfink, Reuter et al. 2021) (≥ 90% amino acid identity and ≥ 90% coverage). Copper and zinc resistance genes were identified using DIAMOND (≥ 90% amino acid identity and ≥ 90% coverage) and the BacMet antibacterial biocide and metal resistance genes database v. 2.0 [15]

### Metagenome-assembled genomes

Metagenomic sequences from conventionally- and pasture-raised pigs were co-assembled separately as well as individually assembled using MEGAHIT v. 1.2.9 [16]. Each sample was mapped back to its respective assembly and the co-assembly using Bowtie and the contigs (≥ 2000 bp) were binned into MAGs using MetaBat 2 v. 2.2.15. The completeness and contamination of both the co-assembled and individually assembled MAGs was calculated with CheckM v1.1.10 [17]. A total of 19,482 MAGs were recovered with greater than 90% completeness and less than 5% contamination. These MAGs were then dereplicated using dRep v.3.2.2 with primary clustering set at 90% and secondary clustering set at 99%, resulting in 1,043 MAGs for subsequent analyses. Taxonomy was then assigned to these 1,043 dereplicated MAGs using GTDB-tk v. 2.1.1 [18] and the GTDB database release 207 [19].

Genes were predicted and annotated using Distilled and Refined Annotation of Metabolism (DRAM) v. 1.3.5 [20] with the dbCAN2 HMMdb release 10.0 and KEGG release 101.0 databases for annotation of CAZymes and other metabolic pathways, respectively. The MAGs were also screened for ARGs using the RGI with the CARD and for copper and zinc resistance genes using DIAMOND (70% identity) and the BacMet antibacterial biocide and metal resistance genes database. A phylogenomic tree was generated from the high quality dereplicated MAGs using PhyloPhlAn v. 3.0 by aligning 400 universal marker genes [21]. The relative abundance of each dereplicated MAG in each sample was determined using CoverM v. 0.6.1 (https://github.com/wwood/CoverM).

All metagenomic sequences and MAGs are publicly available in the National Center for Biotechnology Information’s (NCBI) sequence read archive and genome databases under BioProject PRJNA857725.

### Statistical analysis

Differentially abundant microbial species, ARGs, copper and zinc resistance genes, CAZymes, and MAGs between the conventionally- and pasture-raised pigs were identified with MaAsLin 2 v. 1.10.0 (Mallick et al., 2021) in R 4.2.0. Only those microbial species with an overall percent relative abundance of at least 0.1% within pigs of the same production phase were included in this analysis. For ARGs, copper and zinc resistance genes, and CAZymes, only those features present in at least 25% of the samples were included. Permutational multivariate analysis of variance (PERMANOVA) of the Bray-Curtis dissimilarities was calculated in vegan 2.6-2 to determine the effect of farm type on the structure of the microbial community and resistome. The correlation between the microbiome (species) and the resistome (ARGs) and the microbiome and the CAZyme composition was assessed using Procrustes analysis with the Bray-Curtis dissimilarity NMDS ordinations in vegan.

## RESULTS

### The fecal microbiome of conventionally-vs. pasture-raised pigs

The fecal microbiome of the two groups of pigs differed significantly at the species level (Fig. 1A; PERMANOVA: R^2^ = 0.27, P < 0.001). Among the 141 bacterial species with an overall relative abundance of at least 0.1% in one of the three production phases, 111 were differentially abundant between the conventionally-and pasture-raised pigs (Table S1). *Dialister succinatiphilus, Intestinibaculum porci, Prevotella mizrahii, Pseudoscardovia radai*, and *Sharpea azabuensis* were among the nine bacterial species that were consistently enriched in pasture pigs and *Bacteroides sp*. CAG-709, *Clostridium sp*. CAG-138, *Hallella faecis*, and *Ruminococcus sp*. CAG-177 were relatively more abundant in all pigs raised conventionally (P < 0.05; Fig. 1B). Although *Prevotella copri* was enriched in the conventional growing-finishing pigs, it was the bacterial species with the highest relative abundance overall in both groups of pigs. Other relatively abundant (>0.3%) bacterial species in the gut microbiomes of both conventional and pasture pigs were *Oscillibacter valericigenes* and *Prevotella stercorea* (data not shown). Notably, pasture-raised pigs had significantly greater gut microbial species diversity (Fig. 1C).

**Figure 1.**
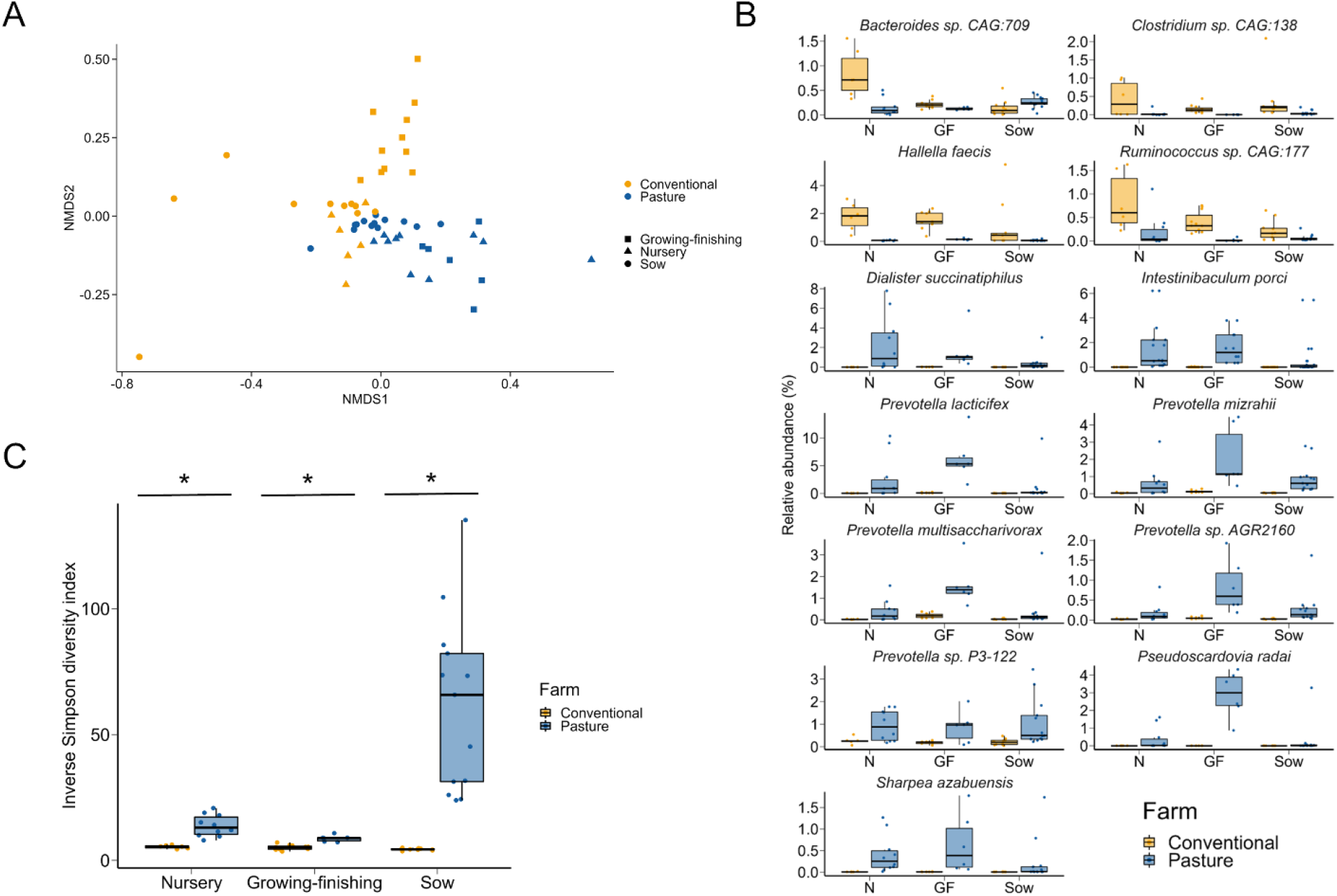
A) Non-metric multidimensional scaling (NMDS) plot of the Bray-Curtis dissimilarities of microbial species in the fecal microbiomes of conventionally- and pasture-raised pigs within three different production phases. B) Percent relative abundance of bacterial species that were differentially abundant in the fecal microbiomes of conventionally- vs. pasture-raised pigs within all three production phases (false discovery rate < 0.05). N = nursery; GF = growing-finishing. C) Inverse Simpson diversity index values for microbial species in the fecal microbiomes of conventionally- and pasture-raised pigs within three different production phases. * = significantly different means (P < 0.05).

CAZymes are enzymes involved in the degradation and synthesis of carbohydrates and are grouped into six different families based on their amino acid sequence similarity: auxiliary activities (AAs), carbohydrate esterases (CEs), carbohydrate-binding modules (CBMs), glycoside hydrolases (GHs), glycosyltransferases (GTs), polysaccharide lyases (PLs) [22]. Here, the CAZyme profiles of the conventionally- and pasture-raised pigs also differed significantly from each other (Fig. 2; PERMANOVA: R^2^ = 0.13, P < 0.001). Of the 209 CAZyme families analyzed, 111 were differentially abundant between the two groups of pigs within one of the production phases. And of those 111 CAZyme families, 14 were differentially abundant within all three production phase groups (Table S2). Within the microbiomes of conventional pigs, there was a consistent enrichment of GH20 (galactosidases), GH31 (glucosidases), and GH68 (levansucrases) families while GH5_2 (endoglucanases), GT111 (galactofuranosyltransferases), and PL1 (pectinases) were relatively more abundant in all pasture-raised pigs (P < 0.05). Other CAZyme families of note that were enriched in the conventionally-raised growing-finishing pigs and sows included GH29 (fucosidases) and GH33 (sialidases), which are associated with mucin degradation. Not surprisingly, the microbiome (species level) was significantly correlated with the CAZyme profiles (Procrustes: R^2^ = 0.62; P = 0.0001)

**Figure 2.**
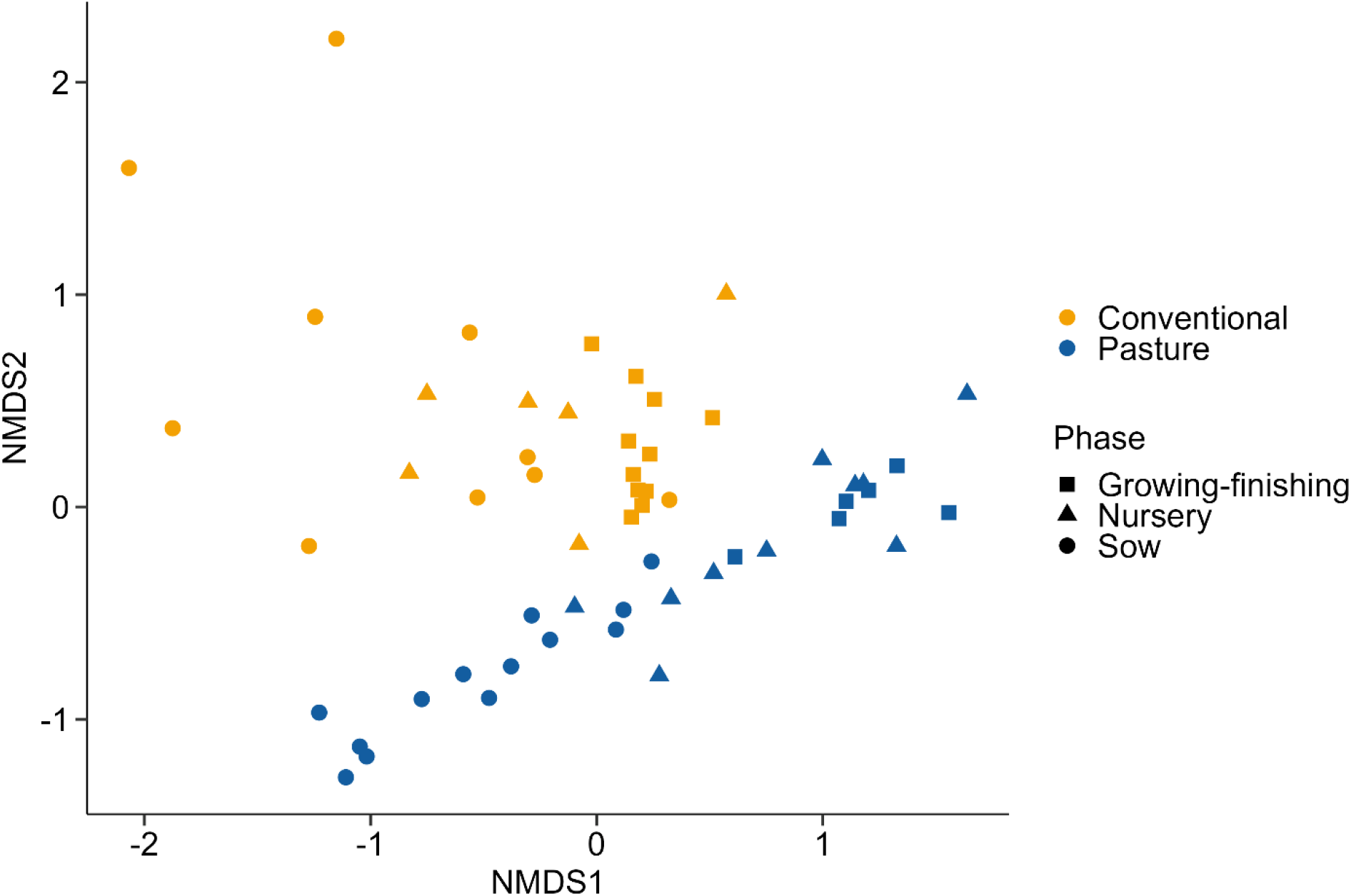
Non-metric multidimensional scaling (NMDS) plot of the Bray-Curtis dissimilarities of carbohydrate-active enzyme (CAZymes) genes in the fecal microbiomes of conventionally- and pasture-raised pigs within three different production phases.

### The resistome of conventionally- vs. pasture-raised pigs

The conventionally- and pasture-raised pigs also had very different resistome profiles based on the relative abundance of ARGs (Fig. 3A; PERMANOVA: R2 = 0.48, P < 0.001). In addition, the relative abundance of nearly all classes of ARGs was significantly higher in the conventional group compared with the pasture pigs (Fig. 3B; Table S3; P < 0.05). There were 271 individual ARGs identified in at least 25% of samples within one of the three production phases and 155 of these ARGs were differentially abundant between the two groups of pigs (Table S4). In addition, of these 155 ARGs, 131 were relatively more abundant in the pigs raised under conventional conditions (P < 0.05). A number of the ARGs consistently enriched in the conventionally-raised pigs were those that are frequently abundant in the pig gut such as *aph(3*’*)- IIIa, erm*(B), *tet*(Q), and *tet*(W) (Fig 4). There was also a significant correlation between the fecal microbiome and resistome (Procrustes: R^2^ = 0.55; P = 0.0001).

**Figure 3.**
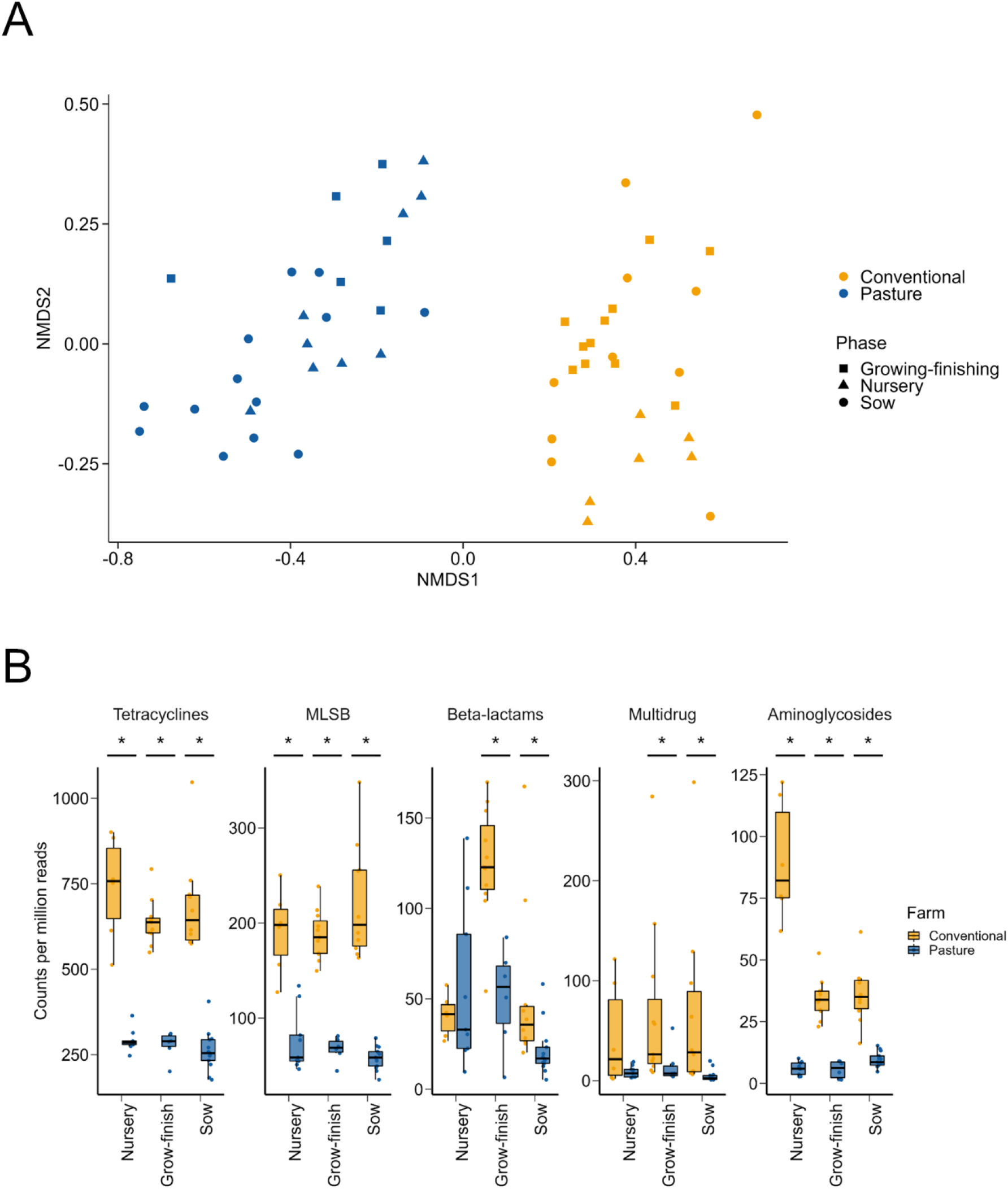
A) Non-metric multidimensional scaling (NMDS) plot of the Bray-Curtis dissimilarities of antimicrobial resistance genes in the fecal microbiomes of conventionally- and pasture-raised pigs within three different production phases. B) Copies per million reads for antimicrobial resistance genes grouped into the antimicrobial classes that they confer resistance to. * = significantly different means (P < 0.05).

**Figure 4.**
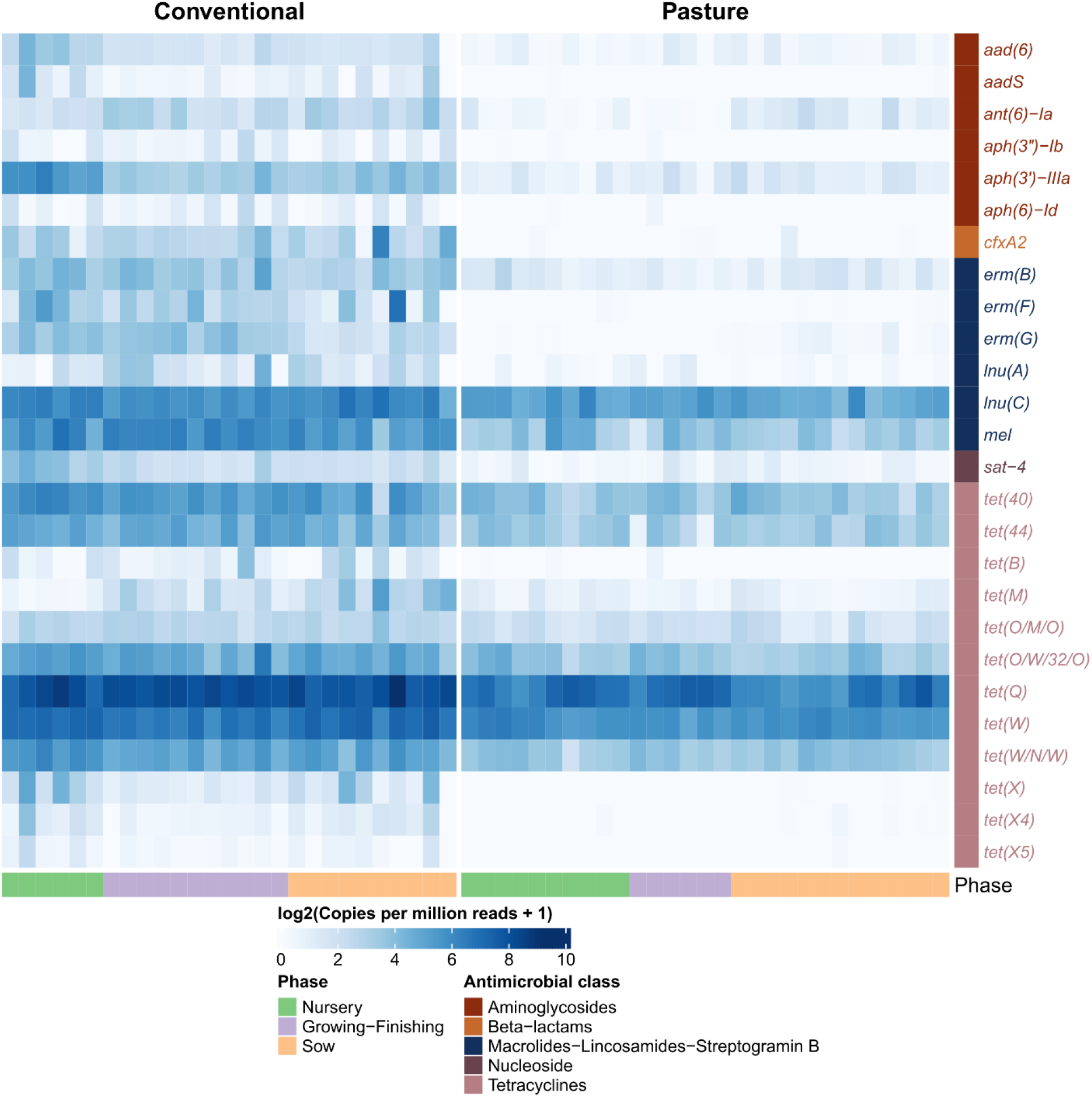
Heat map of the log2 transformed copies per million reads for antimicrobial resistance genes that were differentially abundant in the fecal metagenomes of conventionally- vs. pasture-raised pigs within three different production phases (false discovery rate < 0.05).

Copper and zinc are typically included in pig diets at concentrations that exceed dietary requirements to control bacterial infections and improve growth [23]. However, the inclusion of dietary copper and zinc at these levels may co-select for ARGs located on the same mobile genetic element [24]. Therefore, we also screened the metagenomic sequences for genes associated with copper and zinc resistance. Although there were no copper or zinc resistance genes that were differentially abundant in the microbiomes of growing-finishing pigs, the copper resistance gene *tcrB* was relatively more abundant in the nursery conventional pigs and 28 copper (*cusABCFRS*; *tcrAB*) and zinc resistance genes (e.g. *zitB*) were enriched in the conventionally-raised sows (Table S5).

### Metagenome-assembled genomes

Metagenomes from each group of pigs were assembled and binned into MAGs which were then dereplicated at 99% ANI resulting in 1,043 MAGs with ≥ 90% completeness and ≤ 5% contamination (Fig. 5). These non-redundant MAGs represented 357 genera and 446 species (Table S6). There were 37 and 356 MAGs that could not be assigned to a genus or species, respectively, and thus may represent potentially novel genera and species. Furthermore, only 124 MAGs (11.9%) were assigned to an archaeal or bacterial species with a cultured representative. The most frequently identified bacterial species within the 1,043 MAGs were *Sodaliphilus* sp004557565 (12 MAGs) and *Collinsella* sp002391315 (11 MAGs).

**Figure 5.**
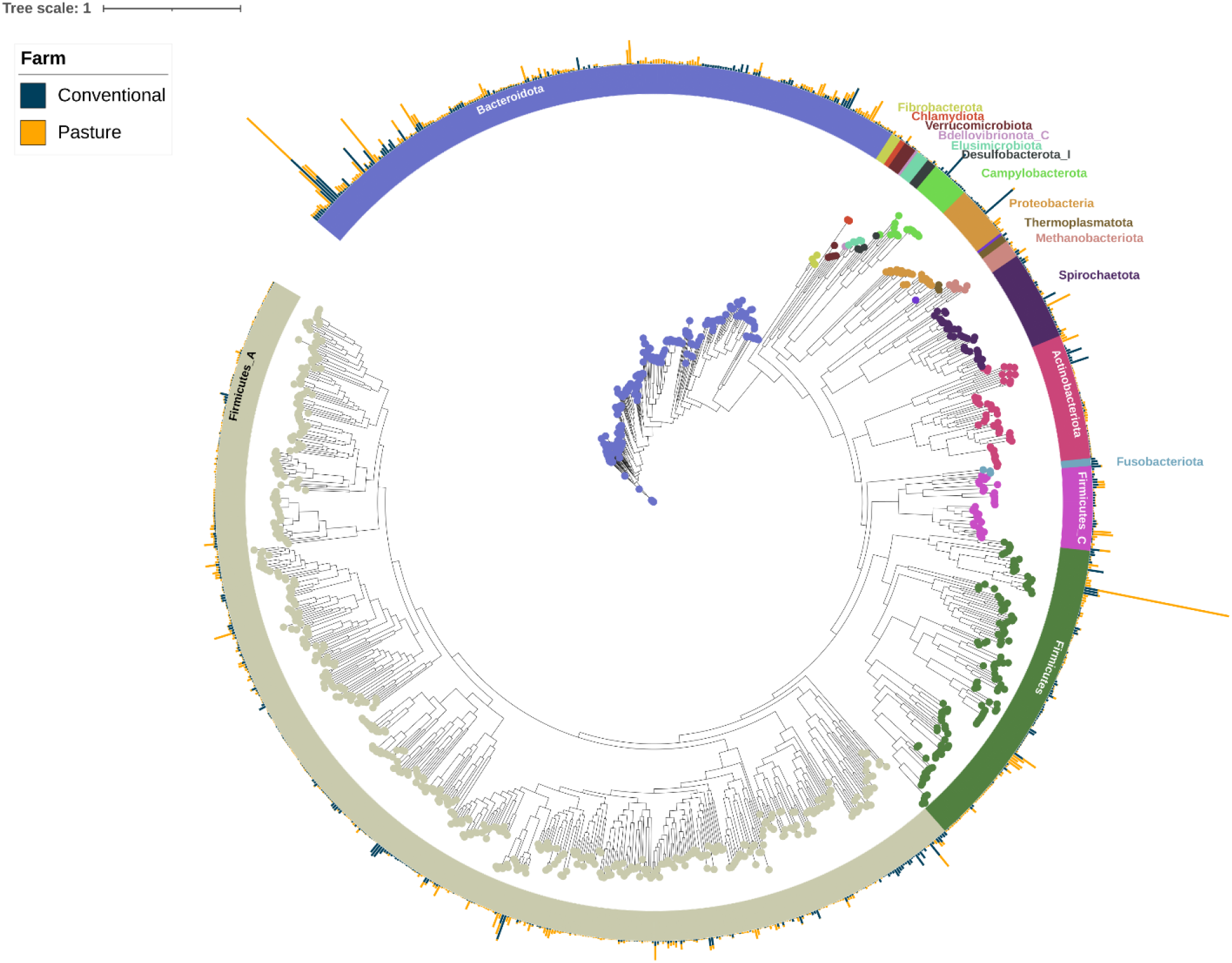
Maximum likelihood phylogenetic tree of the metagenome-assembled genomes (MAGs) based on the alignment of 400 marker genes. MAGs are colored and labeled by GTBD-tk assigned phyla. The outer bars display the percent relative abundance (0% to 3.7%) of each MAG in the conventionally- and pasture-raised pig fecal samples.

Reads from each metagenomic sample were also mapped back to the MAGs with 56.4 ± 0.9% (SEM) of the reads aligning to one of the MAGs (Table S7). There were 35 MAGs that were exclusive to the conventionally-raised pigs including those classified as *Campylobacter coli, Campylobacter hominis*, and *Porphyromonas somerae*. Two *Lactobacillus porci* MAGs were among the 17 MAGs recovered only from the pasture-raised pig fecal metagenomes. There were also two *Pseudoscardovia* spp. MAGs identified in 93.1% of the fecal samples from pasture-raised pigs that were only detected in one conventionally-raised sow. Twelve MAGs were differently abundant between the conventionally- and pasture-raised pigs within all three production phases (Fig. S1). Similar to the unassembled short reads, these MAGs included those classified as *I. porci* and *S. azabuensis* which were relatively more abundant in the pasture pigs and *Limosilactobacillus reuteri, Prevotella* sp002300055, and *Cryptobacteroides* sp000433355 MAGs were enriched in all phases of the conventional pigs. In terms of similarities, three *P. copri* MAGs were among the 10 MAGs with the highest relative abundance in both groups of pigs.

Assembling and binning ARGs is difficult as they are frequently located on mobile genetic elements like plasmids which often have different DNA sequence properties than the host genome [25]. Therefore as expected, far fewer ARGs were detected in the MAGs compared with the unassembled short reads. However, several *tet* genes were identified within the MAGs (Table S8). The most widely distributed *tet* genes were *tet*(Q) (15 MAGs), *tet*(36) (11 MAGs), and *tet*(W/N/W) (8 MAGs). The rRNA methylase genes *erm*(B) and *erm*(X), conferring resistance to macrolides, were detected in a *Streptococcus suis* MAG and a *Corynebacterium* sp. MAG, respectively. Other ARGs of interest included *aph(3*’*)-IIIa* (aminoglycosides) in a *Campylobacter lanienae* MAG, *bla*_OXA-193_ (beta-lactams) in a *Campylobacter coli* MAG, and *cfxA2* (beta-lactams) in two *Sodaliphilus* sp004557565 MAGs. Two *Aerococcus* sp. MAGs that were relatively more abundant in the conventional sow microbiomes were found to be carrying the repressor gene, *tcrY*, from the copper resistance operon *tcrYAZB*, and one of these MAGs also had the *tcrB* gene which encodes a copper resistance efflux ATPase.

Short-chain fatty acids (SCFAs) are produced from non-digestible carbohydrates by certain members of the microbiome in the lower gastrointestinal tract of monogastric animals such as pigs. These SCFAs, which include acetate, butyrate, and propionate, provide energy for the host and have anti-inflammatory properties as well. Here, genes for butyrate production via the butyryl-CoA:acetate CoA-transferase or butyrate kinase pathways were encoded by 84 and 114 MAGs, respectively (Table S9). *Megasphaera elsdenii* and *Sodaliphilus pleomorphus* MAGs carrying genes for butyrate production were also among those significantly associated with conventionally-raised growing-finishing pigs. In addition, several *Fusobacterium* and *Porphyromonas* spp. MAGs enriched in the conventional farm sows encoded genes for one of the two butyrate pathways. Two *Alloprevotella* sp004557185 MAGs with the butyrate kinase genes were relatively more abundant in the pasture-raised nursery phase pigs. Genes for acetate production were identified in the large majority of MAGs (84.7%) while those encoding for propionate were found in only 69 MAGs.

CAZymes involved in the degradation of amorphous cellulose, arabinose, and mixed-linkage glucans were encoded by at least 50% of the MAGs (Table S9). The greatest number of CAZymes and CAZyme families were found in MAGs classified as *Alistipes senegalensis, Parabacteroides faecavium, Phocaeicola plebeius, Phocaeicola vulgatus*, and *Prevotella* sp004554665 (Table S10). There were also 432 CAZyme families detected among all the MAGs, including 164 GH families of which GH2, GH3, GH13, GH23, and GH25 were most prevalent.

## DISCUSSION

The fecal microbiomes and resistomes of pigs raised on a conventional farm were significantly different from those of pigs raised outdoors on pasture. This is not unexpected given the large differences between the two production systems and demonstrates the large impact that management practices can potentially have on the pig gut microbiome and resistome. Pigs on the conventional farm were housed completely indoors and in pens with other pigs as per industry standards whereas the pasture-raised pigs were reared entirely outdoors at a much lower stocking density. Although both groups of pigs were fed a grain-based diet, pigs on pasture will often use their snouts to root in the soil and so it is likely that they are ingesting plants, soil, and other organic matter.

Using both short, unassembled metagenomic reads and MAGs, we identified a number of bacterial species strongly associated with one of the farm types. Some of the bacterial species with a higher relative abundance in the conventionally-raised pigs were those that can be potentially pathogenic in swine such *S. suis* and *C. coli*, although these species are often recovered from healthy pigs as well [26]. Other bacterial species enriched in the conventionally-raised pigs included *L. reuteri, Prevotella* sp002300055, and *Treponema succinifaciens*. Supplementation of swine feed with *L. reuteri* has been linked with improved growth performance and a reduction in certain bacterial pathogens in the pig gut [27-29] and *T. succinifaciens* has also been reported to be associated with the gut microbiome of pigs with higher feed efficiency [30]. *Prevotella* sp002300055 is a placeholder species that we previously identified as abundant in swine gut metagenomes from multiple countries [31].

The gut microbiomes of pigs raised on pasture were enriched with several bacterial species that are not typically abundant in conventionally-raised swine. These included two *Pseudoscardovia* spp. MAGs that were not identified in any of the conventional nursery or growing-finishing pigs and in only one conventional sow. Presently, there are two recognized *Pseudoscardovia* species, *P. radai* and *Pseudoscardovia suis*, both of which were originally characterized using isolates from the gastrointestinal tract of wild boars [32, 33]. Therefore, *Pseudoscardovia* spp., which are closely related to the bifidobacteria, appear to be strongly associated with pigs in the outdoor environment. Two related lactic acid-producing bacterial species, *I. porci* and *S. azabuensis*, were also significantly enriched in the fecal microbiomes of pasture-raised pigs. *S. azabuensis* has been reported to be relatively more abundant in young healthy pigs vs. those with diarrhea [34] and has also been linked with reduced methane emissions in ruminants [35]. *I. porci* is a relatively new bacterial species that was originally isolated from the small intestine of a pig [36] and similar to *S. azabuensis*, it has been associated with lower methane emissions in cattle [37].

Although *P. copri* was enriched in the conventional sow gut microbiome, it was the bacterial species with the highest relative abundance in both conventionally- and pasture-raised pigs and therefore appears to be well-adapted to the domestic pig gut regardless of diet and environment. In pigs, *P. copri* has been associated with the post-weaning phase [4, 38] and fat accumulation [39]. However, *P. copri* is a genetically diverse species that has recently been proposed to contain at least four distinct clades [40] and therefore the *P. copri* strains in the conventionally- and pasture-raised pigs are especially unlikely to be shared given their dissimilar gut microbiomes.

Functionally, the conventional and pasture pig gut microbiomes were also significantly different based on their CAZyme profiles. CAZymes synthesize and degrade carbohydrates and are prevalent in the gut microbiomes of mammals [41]. In gut bacteria, the GHs are the major CAZyme family involved in the breakdown of dietary glycans [42] and here several GH families were differentially abundant between the two pig farms. Notably, certain CAZyme families associated with host mucin degradation such as GH20 (galactosidases) GH29 (fucosidases), and GH33 (sialidases) were enriched in the microbiomes of the conventionally-raised pigs. Bacteria that produce these CAZymes can release oligo- and monosaccharides from mucin glycans found in the gut mucosa which can then be used by other gut bacteria to produce metabolites such as SCFAs through a process called cross-feeding [43].

There were also 19 MAGs enriched in one of the three conventionally-raised pig production groups that carried genes encoding for CAZymes within the GH20, GH29, and GH33 families. Included among these MAGs were two *S. pleomorphus* MAGs and several others from uncultured species such as *Alloprevotella* sp004552155, *Cryptobacteroides* sp000433355, and *Prevotella* sp000434975. *S. pleomorphus* is a relatively new bacterial species within the *Muribaculaceae*, a family associated with mucin degradation [44]. It is possible that the less diverse diet of conventional pigs, compared to that of pigs raised outdoors, favors bacteria that can metabolize host mucins for energy.

Conventionally-raised pigs almost always have a large background of certain antimicrobial-resistant bacteria and ARGs even in the absence of direct antimicrobial exposure [2, 45, 46]. Here, compared to the pasture-raised pigs, the abundance of ARGs within nearly all antimicrobial classes was significantly higher in the conventionally-raised pigs. These ARGs included many that are typically abundant in commercial pigs such as *aph(3*’*)-IIIa, cfxA*2, *erm*(B), *erm*(F), *lnu*(C), *tet*(44), *tet*(Q), *tet*(W), and *tet*(X) [2, 47-49]. Some of these ARGs were also assembled and binned into specific MAGs thus providing taxonomic information. For example, *erm*(B) was linked to a *S. suis* MAG and *tet*(44) was identified in a *Terrisporobacter* sp. MAG, both of which were relatively more abundant in conventionally-raised swine. As mentioned earlier, *S. suis* is a potential pathogen in pigs that can cause arthritis, endocarditis, and meningitis, although it is frequently found in healthy pigs as well [50]. A *C. coli* MAG enriched in the conventionally-raised sows was also carrying *bla*_OXA-193_, a beta-lactamase gene. *C. coli* is prevalent in healthy pigs but has also been implicated in foodborne disease in humans, although less commonly than *Campylobacter jejuni* [51]. The *bla*_OXA-193_ gene has been reported to be relatively common in *C. coli* isolates from multiple sources, including pigs [52].

Although the conventional pigs were not administered any antimicrobials during the study, similar to most conventional swine operations in North America, antimicrobials were used in the facility in the past to promote growth and are still used on a short-term metaphylaxis basis to prevent the spread of disease. Not surprisingly, the relative abundance was also highest for those ARGs that confer resistance to antimicrobials with the longest history of use in swine production such as the beta-lactams, macrolides-lincosamides-streptogramin B (MLS_B_), and tetracyclines. These ARGs appear to be well integrated into the pig gut microbiome without imposing a serious fitness cost to the bacteria that harbour them. Therefore, reducing the abundance of these ARGs likely requires significantly altering the gut microbiome itself as there was a strong association between the species level composition and the resistome here. Conventionally-raised pigs may also be exposed to biocides and disinfectants in their environment as well as high levels of dietary copper and zinc, all of which can co-select for ARGs [53]. Indeed, sows on the conventional farm had a significantly higher relative abundance of a number of different copper and zinc resistance genes, implying there may be some association with the use of these metals at supra-nutritional levels and certain ARGs.

## CONCLUSION

The gut microbiomes of conventionally- and pasture-raised pigs differed significantly from each other with certain bacterial species strongly associated with each group. This included bacterial species such as *Pseudoscardovia* spp. which are rarely detected in conventionally-raised swine but were prevalent in the pasture-raised pigs. Importantly, a large number of ARGs were significantly enriched in the gut microbiome of pigs on the conventional farm. Although reducing antimicrobial resistance in commercial food-producing animals like pigs is very challenging, this study suggests that management practices may potentially make a large difference in this regard.

## Acknowledgements

The authors are greatly appreciative of the owners of the pasture pig farm for allowing access to the pigs and for helping with the sampling. We also thank Cara Service and the animal care staff for their excellent care of the conventionally-raised pigs and for their help with sampling as well. Funding for this project was provided by Agriculture and Agri-Food Canada’s A-Base program.

## References

1. McEwen SA, Collignon PJ. Antimicrobial resistance: a One Health perspective. Microbiol Spectr 2018;6(2):6.2. 10.

2. Pollock J, Muwonge A, Hutchings MR, Mainda G, Bronsvoort BM et al. Resistance to change: AMR gene dynamics on a commercial pig farm with high antimicrobial usage. Sci Rep 2020;10(1):1708.

3. Tunsagool P, Mhuantong W, Tangphatsornruang S, Am-In N, Chuanchuen R et al. Metagenomics of antimicrobial and heavy metal resistance in the cecal microbiome of fattening pigs raised without antibiotics. Appl Environ Microbiol 2021;87(8):e02684–02620.

4. Holman DB, Gzyl KE, Mou KT, Allen HK. Weaning Age and Its Effect on the Development of the Swine Gut Microbiome and Resistome. mSystems 2021;6(6):e0068221.

5. Hobson C, Chan AN, Wright GD. The Antibiotic Resistome: A Guide for the Discovery of Natural Products as Antimicrobial Agents. Chem Rev 2021;121(6):3464–3494.

6. den Besten G, van Eunen K, Groen AK, Venema K, Reijngoud DJ et al. The role of short-chain fatty acids in the interplay between diet, gut microbiota, and host energy metabolism. J Lipid Res 2013;54(9):2325–2340.

7. Bergamaschi M, Tiezzi F, Howard J, Huang YJ, Gray KA et al. Gut microbiome composition differences among breeds impact feed efficiency in swine. Microbiome 2020;8(1):110.

8. Wang X, Tsai T, Deng F, Wei X, Chai J et al. Longitudinal investigation of the swine gut microbiome from birth to market reveals stage and growth performance associated bacteria. Microbiome 2019;7(1):109.

9. Chen S, Zhou Y, Chen Y, Gu J. fastp: an ultra-fast all-in-one FASTQ preprocessor. Bioinformatics 2018;34(17):i884–i890.

10. Langmead B, Salzberg SL. Fast gapped-read alignment with Bowtie 2. Nat Methods 2012;9(4):357–359.

11. Li H, Handsaker B, Wysoker A, Fennell T, Ruan J et al. The Sequence Alignment/Map format and SAMtools. Bioinformatics 2009;25(16):2078–2079.

12. Quinlan AR, Hall IM. BEDTools: a flexible suite of utilities for comparing genomic features. Bioinformatics 2010;26(6):841–842.

13. Alcock BP, Raphenya AR, Lau TTY, Tsang KK, Bouchard M et al. CARD 2020: antibiotic resistome surveillance with the comprehensive antibiotic resistance database. Nucleic Acids Res 2020;48(D1):D517–D525.

14. Clausen P, Aarestrup FM, Lund O. Rapid and precise alignment of raw reads against redundant databases with KMA. BMC Bioinformatics 2018;19(1):307.

15. Pal C, Bengtsson-Palme J, Rensing C, Kristiansson E, Larsson DG. BacMet: antibacterial biocide and metal resistance genes database. Nucleic Acids Res 2014;42(Database issue):D737–743.

16. Li D, Liu CM, Luo R, Sadakane K, Lam TW. MEGAHIT: an ultra-fast single-node solution for large and complex metagenomics assembly via succinct de Bruijn graph. Bioinformatics 2015;31(10):1674–1676.

17. Parks DH, Imelfort M, Skennerton CT, Hugenholtz P, Tyson GW. CheckM: assessing the quality of microbial genomes recovered from isolates, single cells, and metagenomes. Genome Res 2015;25(7):1043–1055.

18. Chaumeil PA, Mussig AJ, Hugenholtz P, Parks DH. GTDB-Tk v2: memory friendly classification with the genome taxonomy database. Bioinformatics 2022;38(23):5315–5316.

19. Parks DH, Chuvochina M, Rinke C, Mussig AJ, Chaumeil PA et al. GTDB: an ongoing census of bacterial and archaeal diversity through a phylogenetically consistent, rank normalized and complete genome-based taxonomy. Nucleic Acids Res 2022;50(D1):D785–D794.

20. Shaffer M, Borton MA, McGivern BB, Zayed AA, La Rosa SL et al. DRAM for distilling microbial metabolism to automate the curation of microbiome function. Nucleic Acids Res 2020;48(16):8883–8900.

21. Asnicar F, Thomas AM, Beghini F, Mengoni C, Manara S et al. Precise phylogenetic analysis of microbial isolates and genomes from metagenomes using PhyloPhlAn 3.0. Nat Commun 2020;11(1):2500.

22. Drula E, Garron ML, Dogan S, Lombard V, Henrissat B et al. The carbohydrate-active enzyme database: functions and literature. Nucleic Acids Res 2022;50(D1):D571–D577.

23. Willing BP, Pepin DM, Marcolla CS, Forgie AJ, Diether NE et al. Bacterial resistance to antibiotic alternatives: a wolf in sheep’s clothing? Anim Front 2018;8(2):39–47.

24. Baker-Austin C, Wright MS, Stepanauskas R, McArthur JV. Co-selection of antibiotic and metal resistance. Trends Microbiol 2006;14(4):176–182.

25. Maguire F, Jia B, Gray KL, Lau WYV, Beiko RG et al. Metagenome-assembled genome binning methods with short reads disproportionately fail for plasmids and genomic Islands. Microb Genom 2020;6(10).

26. Votsch D, Willenborg M, Weldearegay YB, Valentin-Weigand P. Streptococcus suis - The “two faces” of a pathobiont in the porcine respiratory tract. Front Microbiol 2018;9:480.

27. Wang W, Zijlstra RT, Ganzle MG. Feeding Limosilactobacillus fermentum K9-2 and Lacticaseibacillus casei K9-1, or Limosilactobacillus reuteri TMW1.656 reduces pathogen load in weanling pigs. Front Microbiol 2020;11:608293.

28. Valeriano VD, Balolong MP, Kang DK. Probiotic roles of Lactobacillus sp. in swine: insights from gut microbiota. J Appl Microbiol 2017;122(3):554–567.

29. Wang G, Wang X, Ma Y, Cai S, Yang L et al. Lactobacillus reuteri improves the development and maturation of fecal microbiota in piglets through mother-to-infant microbe and metabolite vertical transmission. Microbiome 2022;10(1):211.

30. Quan J, Wu Z, Ye Y, Peng L, Wu J et al. Metagenomic characterization of intestinal regions in pigs with contrasting feed efficiency. Front Microbiol 2020;11:32.

31. Holman DB, Kommadath A, Tingley JP, Abbott DW. Novel Insights into the Pig Gut Microbiome Using Metagenome-Assembled Genomes. Microbiol Spectr 2022;10(4):e0238022.

32. Killer J, Havlik J, Bunesova V, Vlkova E, Benada O. Pseudoscardovia radai sp. nov., another representative of a new genus within the family Bifidobacteriaceae isolated from the digestive tract of a wild pig (Sus scrofa scrofa). Int J Syst Evol Microbiol 2014;64(9):2932–2938.

33. Killer J, Mrazek J, Bunesova V, Havlik J, Koppova I et al. Pseudoscardovia suis gen. nov., sp. nov., a new member of the family Bifidobacteriaceae isolated from the digestive tract of wild pigs (Sus scrofa). Syst Appl Microbiol 2013;36(1):11–16.

34. Yang Q, Huang X, Zhao S, Sun W, Yan Z et al. Structure and function of the fecal microbiota in diarrheic neonatal piglets. Front Microbiol 2017;8:502.

35. Stewart RD, Auffret MD, Warr A, Walker AW, Roehe R et al. Compendium of 4,941 rumen metagenome-assembled genomes for rumen microbiome biology and enzyme discovery. Nat Biotechnol 2019;37(8):953–961.

36. Kim JS, Choe H, Lee YR, Kim KM, Park DS. Intestinibaculum porci gen. nov., sp. nov., a new member of the family Erysipelotrichaceae isolated from the small intestine of a swine. J Microbiol 2019;57(5):381–387.

37. Smith PE, Kelly AK, Kenny DA, Waters SM. Differences in the composition of the rumen microbiota of finishing beef cattle divergently ranked for residual methane emissions. Front Microbiol 2022;13:855565.

38. Gaio D, DeMaere MZ, Anantanawat K, Chapman TA, Djordjevic SP et al. Post-weaning shifts in microbiome composition and metabolism revealed by over 25 000 pig gut metagenome-assembled genomes. Microb Genom 2021;7(8).

39. Chen C, Fang S, Wei H, He M, Fu H et al. Prevotella copri increases fat accumulation in pigs fed with formula diets. Microbiome 2021;9(1):175.

40. Tett A, Huang KD, Asnicar F, Fehlner-Peach H, Pasolli E et al. The Prevotella copri complex comprises four distinct clades underrepresented in westernized populations. Cell Host Microbe 2019;26(5):666–679 e667.

41. Wardman JF, Bains RK, Rahfeld P, Withers SG. Carbohydrate-active enzymes (CAZymes) in the gut microbiome. Nat Rev Microbiol 2022;20(9):542–556.

42. Kaoutari AE, Armougom F, Gordon JI, Raoult D, Henrissat BJNRM. The abundance and variety of carbohydrate-active enzymes in the human gut microbiota. 2013;11(7):497–504.

43. Berkhout MD, Plugge CM, Belzer CJG. How microbial glycosyl hydrolase activity in the gut mucosa initiates microbial cross-feeding. 2022;32(3):182–200.

44. Pereira FC, Wasmund K, Cobankovic I, Jehmlich N, Herbold CW et al. Rational design of a microbial consortium of mucosal sugar utilizers reduces Clostridiodes difficile colonization. Nat Commun 2020;11(1):5104.

45. Holman DB, Chenier MR. Impact of subtherapeutic administration of tylosin and chlortetracycline on antimicrobial resistance in farrow-to-finish swine. FEMS Microbiol Ecol 2013;85(1):1–13.

46. Muurinen J, Richert J, Wickware CL, Richert B, Johnson TA. Swine growth promotion with antibiotics or alternatives can increase antibiotic resistance gene mobility potential. Sci Rep 2021;11(1):5485.

47. Gaire TN, Odland C, Zhang B, Ray T, Doster E et al. The impacts of viral infection and subsequent antimicrobials on the microbiome-resistome of growing pigs. Microbiome 2022;10(1):118.

48. Joyce A, McCarthy CGP, Murphy S, Walsh F. Antibiotic resistomes of healthy pig faecal metagenomes. Microb Genom 2019;5(5).

49. Munk P, Knudsen BE, Lukjancenko O, Duarte ASR, Van Gompel L et al. Abundance and diversity of the faecal resistome in slaughter pigs and broilers in nine European countries. Nat Microbiol 2018;3(8):898–908.

50. Goyette-Desjardins G, Auger JP, Xu J, Segura M, Gottschalk M. Streptococcus suis, an important pig pathogen and emerging zoonotic agent-an update on the worldwide distribution based on serotyping and sequence typing. Emerg Microbes Infect 2014;3(6):e45.

51. Patrick ME, Henao OL, Robinson T, Geissler AL, Cronquist A et al. Features of illnesses caused by five species of Campylobacter, Foodborne Diseases Active Surveillance Network (FoodNet) - 2010-2015. Epidemiol Infect 2018;146(1):1–10.

52. Zhang P, Zhang X, Liu Y, Cui Q, Qin X et al. Genomic insights into the increased occurrence of campylobacteriosis caused by antimicrobial-resistant Campylobacter coli. mBio 2022;13(6):e0283522.

53. Li X, Rensing C, Vestergaard G, Arumugam M, Nesme J et al. Metagenomic evidence for co-occurrence of antibiotic, biocide and metal resistance genes in pigs. Environ Int 2022;158:106899.

